# Dorsal skin biopsies: A non-lethal sampling method for studying amphibians, including the highly endangered Harlequin frogs (Bufonidae: *Atelopus*)

**DOI:** 10.64898/2026.02.04.703782

**Authors:** María José Navarrete Méndez, Amanda B. Quezada Riera, Andrea Terán-Valdez, Elena Naydenova, Luis A. Coloma, Rebecca D. Tarvin

## Abstract

Non-lethal sampling methods are increasingly essential for amphibian research as global declines intensify and many species persist in small, vulnerable populations. Skin biopsies offer a promising alternative to whole-animal collection and other minimally invasive approaches; however, systematic evaluations of recovery and impacts on body condition remain limited. Here, we assess the effects of small (2-mm) dorsal skin biopsies in four frog species, including three highly endangered Harlequin frogs (*Atelopus bomolochos*, *A. balios*, *A. longirostris*) and the Gualataco marsupial frog (*Gastrotheca riobambae*). Under controlled laboratory conditions and in semi-natural enclosures, we monitored wound healing, survival, and body mass trajectories in biopsied and control individuals over a one-month period. Across all species, biopsy sites fully healed within approximately three weeks, following consistent stages of re-epithelialization and subsequent repigmentation. No biopsy-related mortality was observed, and body mass did not differ between biopsied and control individuals, indicating no detectable effects of skin biopsies on body condition during the wound-healing period. Occasional minor post-biopsy reactions resolved without intervention within the observation period. We additionally report anecdotal field recovery observations for three other species (*A. coynei*, *A. laetissimus*, and *A.* sp. *aff. longirostris*), indicating survival and visible wound closure following release. Together, these results indicate that small dorsal skin biopsies represent a safe, non-lethal sampling method for amphibians, including highly endangered taxa. By providing sufficient tissue for diverse downstream applications—such as chemical analyses, genomics, transcriptomics, microbiome characterization, and disease detection—this approach expands the range of questions that can be addressed while minimizing harm to threatened species.

---

Amphibians hold many secrets about the mechanisms that generate and sustain biological diversity. Over millions of years, they have colonized nearly every continent and an extraordinary range of ecosystems, including some of the planet’s most extreme environments (Duellman and Trueb 1994; Carvalho et al. 2010; Hopkins and Brodie 2015). For example, some species inhabit the highest elevations on Earth, such as the Himalayas (Yang et al. 2019; Wang et al. 2018) and the Andes (Navas 1997), where they endure extreme cold and low oxygen levels. Others live at sea level under high salinity conditions comparable to seawater (Hopkins and Brodie 2015), or in deserts and arid regions, where they withstand high temperatures and limited water availability (Carvalho et al. 2010). More recently, some amphibians have begun to adapt to anthropogenic pressures, surviving in highly polluted environments and enduring contaminants such as pesticides (Hua et al. 2015). As such, amphibians represent an ideal system for studying adaptive responses, phenotypic plasticity, and uncovering global patterns in ecology and biodiversity (Womack et al. 2022).

The study of amphibians is not only intellectually compelling but also critically urgent, as they are experiencing dramatic global population declines, with an estimated 41% of extant species threatened with extinction (IUCN Red List of Threatened Species, 2025) and many already presumed extinct (Stuart et al. 2004; Womack et al. 2022). Research can provide critical tools to expand our understanding of amphibians and inform more effective conservation strategies aimed at slowing or reversing declines (Moor et al. 2022; Park and Do 2023). At a minimum, such studies allow us to document unique traits before they disappear (Tyler et al. 1983; Yotsu et al. 1990). Yet many research questions require extensive sampling—across both numbers of individuals and geographic ranges—to identify patterns with statistical confidence and evolutionary relevance. While this was historically feasible given the former abundance of amphibians, it is now virtually impossible for some species and possibly detrimental for others. For instance, more than 300 individuals of *Atelopus chiriquiensis* and over 400 of *Atelopus varius* (Fuhrman et al. 1976; Kim et al. 1975) were sampled in the 1970s to characterize the presence of chemical defenses in these species, likely at a time when they were highly abundant. However, both species experienced drastic population declines in the late 1980s and 1990s, along with many other *Atelopus* species, and are now listed Extinct and Critically Endangered, respectively (IUCN Red List of Threatened Species, n.d.).

Given such challenges, the amphibian research community must increasingly adopt and refine alternative, non-lethal sampling methods that allow for rigorous investigation without further endangering vulnerable populations. Common approaches that have been used include toe clipping and buccal, cloacal, or skin swabbing (Zemanova et al. 2025). These methods have been widely applied in genetic studies and disease detection (Zemanova et al. 2025), while toe clipping has also been used in ecological research to mark individuals (Perry et al. 2011) and estimate age (Peng et al. 2022). However, swabs can yield limited amounts of DNA (Ringler 2018; Rainey et al. 2024); but see (Ambu and Dufresnes 2023) and both swabbing and toe clipping restrict the scope of downstream analyses, particularly those requiring information on skin structure, composition, or physiology. For instance, chemical defenses cannot be reliably assessed using these techniques, especially when toxins are present at low concentrations or when swabs are not paired with stimulation methods that promote toxin release.

A promising alternative is the use of skin biopsies. This method employs a small, sharp-ended biopsy tool, the same used for humans, to remove a portion of full-thickness skin tissue including epidermis, dermis, and in some cases, the hypodermis, too. In many amphibian species, there are regions of the skin that are not tightly attached to the underlying musculature, allowing the skin to readily separate away from the body and the collection of tissue with minimal invasiveness. Skin biopsies have proven successful to detect and quantify the potent neurotoxin tetrodotoxin (TTX) in newts of the genus *Taricha (Bucciarelli et al. 2014; Vaelli et al. 2020; Hanifin et al. 2002)*, to assess the presence of reactive oxygen species in frog skin for eco-monitoring of polluted environments (D’Errico et al. 2018), and to evaluate skin health in the Hellbender salamander (*Cryptobranchus alleganiensis*) (Hardman 2020). Key advantages of this approach include eliminating the need to collect or sacrifice whole animals, minimizing physical harm, and avoiding disruption of reproductive behavior or the removal of individuals from their natural habitats (Bucciarelli et al. 2014).

However, there is currently no systematic documentation of the recovery of animals sampled using skin biopsies, which limits our ability to evaluate the method’s effectiveness in minimizing harm and assess its potential impacts on body condition and post-release survival. In the present study, we address this gap by documenting wound healing, weight change, and survival following biopsy sampling in three amphibian species representing two families. Specifically, we examine two species of the highly endangered *Atelopus* frogs (Bufonidae) (Lötters et al. 2023; Coloma and Duellman 2025b) in laboratory conditions, as well as in *Gastrotheca riobambae* (Hemiphractidae) both in the laboratory and in semi-natural enclosures. We also report field observations of individuals from other *Atelopus* species after release, providing valuable insights into both the short- and long-term outcomes of this non-lethal technique. Our findings offer important guidance on skin biopsies as an alternative sampling method that enables data collection while minimizing the detrimental effects of sampling on already vulnerable populations, supporting the development of ethical and effective practices in amphibian research and conservation of species with small population sizes.

## Materials and Methods

### Ethics statement

Animal procedures in this study were conducted in accordance with the Institutional Animal Care and Use Committee (IACUC) of the University of California, Berkeley (AUP-2019-08-12457-1).

### Species sampling and experimental design

The experiment was conducted from May 1 to June 1, 2024, under controlled laboratory conditions at the Centro Jambatu de Investigación y Conservación de Anfibios (CJ) in San Rafael, Pichincha, Ecuador. We used laboratory-bred individuals from three *Atelopus* species successfully reproduced ex-situ: *A. bomolochos*, *A. balios*, and *A. longirostris*. In addition, we included individuals of *Gastrotheca riobambae* to evaluate the methodology in a different taxon and to assess recovery in a semi-enclosed setting. *G. riobambae* is native to Quito, Pichincha, Ecuador, and has disappeared from much of its historic range. As part of the ex situ breeding and ongoing reintroduction efforts of CJ, some individuals are maintained in outdoor seminatural-enclosures, offering the opportunity to compare recovery between frogs housed under terrarium-controlled conditions and those exposed to more natural environmental conditions.

The species included in this study differ markedly in conservation status, geographic range, ecological breadth, and body size. The three *Atelopus* species examined are all narrowly distributed Andean endemics occupying distinct elevational and forest types across Ecuador, and they span a range of adult body sizes as measured by snout-vent length (SVL). *A. bomolochos* is a moderately sized species that occurs in the highland montane forest in the southeastern of the Andes, and is the largest *Atelopus* included in this study, with adult males averaging 39.6 mm SVL (range: 38.4–40.8 mm; n = 2) and adult females averaging 47.5 mm SVL (range: 43.9–51.0 mm; n = 5) (Ron et al. 2024; (Coloma and Duellman 2025b). *A. balios* inhabits montane foothill forests of the southern Pacific lowlands and is a relatively small-bodied species, with adult males averaging 27.8 mm SVL (range: 27.1–29.9 mm; n = 5) and adult females averaging 37.1 mm SVL (range: 30.3 – 39.0 mm; n = 8) (Ron et al. 2024; (Coloma and Duellman 2025b)*. A. longirostris* is known from the western slopes of Cordillera Occidental and is a moderate-sized species, with adult males averaging 32.7 mm (range: 30.3 – 35.1 mm; n = 5) and adult females averaging 42.5 mm (range: 40.7 – 47.1 mm; n = 10) (Coloma and Duellman 2025b)Ron et al. 2024; (Coloma and Duellman 2025b). *A. bomolochos, A. balios,* and *A. longirostris* are listed as Critically Endangered on the IUCN Red List of Threatened Species ((IUCN Red List of Threatened Species, n.d.)18, 2019, 2022). By contrast, *G. riobambae* has a substantially broader geographic and environmental distribution. It inhabits humid montane shrubs and grasslands, also dry rocky slopes in montane forest, as well as forests and inter-Andean valleys of northern and central Ecuador. This species is larger than any of the *Atelopus* species sampled, with adult males averaging 43.0 mm SVL (range: 34.1–56.8 mm; n = 81) and adult females averaging 48.6 mm SVL (range: 33.3–66.4 mm; n = 106) (Ron et al. 2024, (Coloma and Duellman 2025a), and is currently categorized as Vulnerable by the IUCN (IUCN Red List of Threatened Species, n.d.) and Endangered by Coloma and Duellman (2025b).

A total of 10 individuals per *Atelopus* species were included in the study, with five undergoing the biopsy procedure and five serving as controls to monitor survival and assess if body weight (a proxy for body condition) remained comparable between treatments. For *G. riobambae*, we included 20 individuals, with 10 held in laboratory-terrarium conditions and 10 in semi-controlled enclosures. Within each housing condition, five individuals were sampled using skin biopsies and five served as unmanipulated controls. Each individual’s body weight and snout–vent length (SVL) were recorded using a scale accurate to 0.01 grams and a caliper accurate to 0.1 millimeters. Additionally, individuals were photo-identified (dorsal color patterns) using a Canon 6D camera with a Sigma 105 mm macro lens.

All *Atelopus* individuals were housed in groups in tanks measuring 30 x 22.5 x 16 cm and *Gastrotheca riobambae* individuals in tanks measuring 43 x 32 x 28 cm for the duration of the monitoring period, with each treatment group maintained separately. Tanks were pre-washed and disinfected prior to use. Each was lined with moistened paper towels, commonly used in veterinary settings, and included autoclaved dry leaves to provide shelter. Environmental conditions were maintained to resemble the natural habitat of each species: *A. bomolochos* was kept at a mean temperature of 21.5 °C (range: 20.7–23.5 °C), while *A. balios*, *A. longirostris*, and *G. riobambae* were maintained at 24.5 °C (range: 23.0–25.2 °C), all under a 12-hour light/12-hour dark photoperiod. The semi-enclosed outdoor population of *G. riobambae* was kept in their original tanks with access to water and shade, under ambient temperature and humidity conditions (maximum temperature = 21.2 °C, minimum temperature = 9.7 °C; relative humidity ≈ 50%; Empresa Pública Metropolitana de Agua Potable y Saneamiento de Quito [EPMAPS] 2026). During the monitoring period, all individuals were periodically fed crickets *Gryllus* sp. (*assimilis* complex) (Orthoptera), flies *Drosophila melanogaster,* and larvae of the fly *Megaselia scalaris* (Diptera). No acclimatization period was established prior to the experiment, as all individuals had been raised in the same facility.

To document recovery, a thorough assessment of wound healing and overall animal health was conducted by CJ’s veterinarian; individuals were monitored for signs of distress, including inability to jump, failure to right themselves when placed on their backs, or any anomalous behavior. In addition, body weight measurements and photographic documentation was conducted every three days. Animals were handled with fresh nitrile gloves at all times between treatments and species to minimize the risk of bacterial or fungal cross-contamination. Monitoring continued for one week following complete wound healing in all sampled individuals (by May 25, 2024), and the experiment concluded on June 1, 2024. Wound healing was defined in a functional sense as complete re-epithelialization, with no visible open wounds or signs of infection (see Fig. 5 in Ishii et al. 2021 for a schematic of amphibian skin wound healing).

### Biopsy procedure

This non-lethal biopsy technique was initially proposed for the study of chemical defenses in newts by (Hanifin et al. 1999), with subsequent modifications introduced by (Bucciarelli et al. 2014). However, detailed procedural guidance for its application has not been fully described. Here, we provide a comprehensive, step-by-step description of the biopsy procedure, accompanied by images (Fig. 1A) and a protocol video illustrating hand placement, animal positioning, and biopsy execution. All handling was conducted using disposable nitrile gloves, which were replaced between individuals to minimize the risk of pathogen transmission. For capture and restraint, *Atelopus* and *Gastrotheca* individuals were gently secured by encircling the waist and holding the femur with the non-dominant hand.

**Figure 1.**
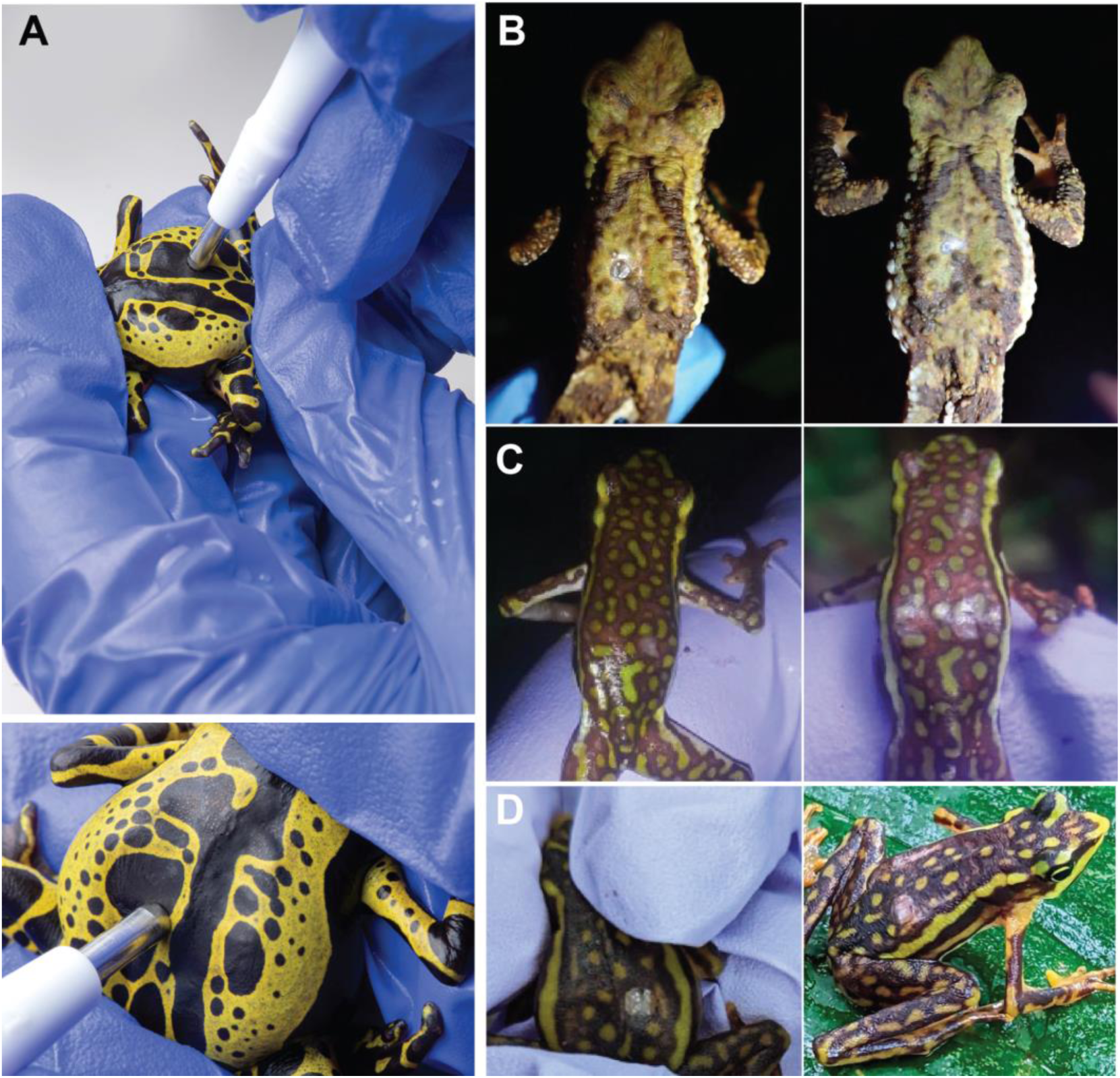
Skin biopsy methodology and post-biopsy recovery under field conditions. **A.** Skin biopsy procedure. The individual is restrained using the non-dominant hand: the middle finger provides ventral support, the index finger stabilizes the head, and the thumb applies gentle pressure to the posterior portion of the body to maintain the dorsal skin taut. A sterile biopsy punch is applied perpendicular to the dorsum and rotated 360° in both directions to fully excise the tissue. The amount of pressure applied was adjusted according to skin thickness and body size, with greater pressure required for larger species with thicker skin; however, pressure was assessed on a case-by-case basis. Images: José Vieira. **B.** Field documentation of recovery in *Atelopus laetissimus* from the Sierra Nevada de Santa Marta, Colombia, obtained during ongoing population monitoring. The individual was biopsied on July 8, 2024; photographs correspond to July 21 (left) and August 8, 2024 (right). Images: Tobias Hildwein. **C.** Post-biopsy recovery in field-sampled *A. coynei.* The individual was biopsied on January 1, 2023; photographs correspond to January 9 (left) and January 14, 2023 (right), showing progressive re-epithelialization of the biopsy site. Images: Kléver Vélez. **D.** Post-biopsy recovery in field-sampled *A.* sp. (*aff. longirostris*). The individual was biopsied on December 26, 2022; photographs were taken on January 5 (left) and January 14, 2023 (right), documenting wound closure and tissue regeneration. Images: Kléver Vélez.

Tissue samples were collected from the dorsal surface of each specimen using a sterile, disposable 2 mm skin biopsy punch (Miltex™, Integra™, available through Fisher Scientific). Chemical anesthesia was not used prior to sampling. In small-bodied animals, inappropriate dosing of anesthetics can lead to adverse effects on the central nervous and cardiovascular systems (Lemke and Dawson 2000; Chatigny et al. 2017). Moreover, there is limited experimental data on the use of local anesthetics in amphibians, and appropriate dosage guidelines are lacking (Chatigny et al. 2017). However, to minimize discomfort, ice was applied to the selected biopsy site prior to sampling to cold-numb and desensitize the area.

To facilitate tissue collection, the frog’s body was gently flexed to stretch the dorsal skin, allowing the biopsy tool to cut more efficiently. Individuals were restrained using the middle finger, index finger, and thumb of the non-dominant hand: the middle finger provided ventral support, the index finger stabilized the head, and the thumb applied gentle pressure to the posterior portion of the body. This positioning maintained the skin taut, producing a cleaner and more precise biopsy, as the punch performs optimally on stretched tissue (Fig. 1A). Finger placement and applied pressure were adjusted according to individual body size. Using the dominant hand, the biopsy punch was applied perpendicular to the skin surface at the selected site (Fig. 1A). Gentle pressure was applied while rotating the tool 360° in both directions to fully excise the tissue. In cases where tissue was not completely detached due to insufficient pressure during punching, fine forceps and small sterilized surgical scissors were used to carefully complete removal.

Additional considerations included skin thickness and the selection of body regions with sufficiently thick skin, which provide greater protection to the underlying musculature and internal organs during sampling. Preference should be given to areas where the skin is relatively loose and not tightly adherent to the underlying bone or musculature. In *Atelopus*, such regions are typically located away from the cranium, whereas tighter skin attachment near the skull limits biopsy placement in some individuals. This close attachment, with little to no intervening tissue between the cranium and the skin, was particularly evident in *A. balios* and *A. longirostris*, which have a more slender body condition than *A. bomolochos* and especially than *G. riobambae*. Accordingly, biopsies were taken from a standardized site on the right dorsolateral surface at the midpoint of the dorsum, lateral to the vertebral column and anterior to the pelvic girdle.

### Field settings

We also monitored the recovery of animals under wild conditions in two field environments. In the first, two individuals of *Atelopus coynei* and *Atelopus* sp. (*aff*. *longirostris*) were monitored following biopsy sampling conducted on January 1, 2023 and December 26, 2022, respectively in the locality of San Jacinto, cantón Mira, Carchi province. Each individual was encountered multiple times after sampling (four times for *A. coynei* and five times for *Atelopus* sp. (*aff. longirostris*)). However, only photographs representing mid-recovery and the final encounter prior to the completion of surveys are shown, as photographs were not standarized across encounters and recovery monitoring was not originally planned as part of this study. For *A. coynei*, the images correspond to January 9 and January 14, 2023 (8 and 13 days post biopsy), while for *A.* sp. (*aff. longirostris)*, photographs were taken on January 5 and January 14, 2023 (10 and 19 days post biopsy) (Fig. 1C–D).

In the second, as part of a separate monitoring project, we made opportunistic observations of released individuals of *A. laetissimus* in the Sierra Nevada de Santa Marta, Colombia after skin biopsies were taken. Specifically, one *A. laetissimus* individual was photographed on two occasions following a dorsal skin biopsy collected on July 8, 2024: the first approximately two weeks later, on July 21, and the second about one month post-biopsy, on August 8, 2024 (Fig. 1B). Due to limited access to the sampling sites, we were unable to include a large number of individuals in either study and thus could not perform statistics on field-sampled animals.

### Statistics and recovery assessment

All statistical analyses were conducted in Rstudio version 2025.5.0.496 (Posit Team 2025). To assess whether biopsy sampling affected body condition, we compared body mass between control and treatment groups using two approaches: (1) final body mass, measured on June 1, 2024, and (2) longitudinal body mass measurements collected every three days throughout the experiment (May 1–June 1, 2024). Analyses were performed separately for each species. For *Gastrotheca riobambae*, individuals from laboratory conditions and semi-natural enclosures were analyzed both jointly and separately to evaluate consistency across housing conditions. For the analysis of final body mass, we evaluated the normality of mass (g) distributions within each species using the Shapiro–Wilk test. As all distributions met the assumption of normality (p > 0.05), Welch’s t-tests were used to compare final masses between treatment and control groups within each species. Two individuals from the control group—one *A. balios* and one *A. bomolochos*—died during the third week of the experiment, and their data were excluded from the analysis (see Results).

To analyze mass trajectories over time, we fitted a linear mixed-effects model using the *nlme* package, with body mass as the response variable. The model included initial mass as a covariate to control for baseline differences among individuals. Fixed effects included initial mass, treatment group (control vs. biopsy), and sampling time point, defined as the date of measurement following the biopsy procedure and treated as a categorical variable because measurements were collected at regular three-day intervals from May 1 to June 1, 2024. Individual identity was included as a random effect to account for repeated measurements over time. Because species differed substantially in body size, analyses were conducted separately for each species. Model residuals were visually inspected and assessed for normality using a Shapiro–Wilk test on Pearson residuals.

## RESULTS

### Mortality

Two individuals from the control groups—*Atelopus balios* (AQCT03) and *A. bomolochos* (AQCT07)—died on May 30. No signs of injury or lacerations associated with chytrid fungus were observed and no chytrid spores were found under microscopic analysis of shedding skin, making it difficult to determine the exact cause of death. According to the veterinarian and CJ staff involved in the study, the deaths may have been related to changes in experimental handling, as both individuals had previously been housed in larger terrariums under different environmental conditions than those of the experimental tanks.

### Healing and recovery process

The healing of the incision was monitored every three days by recording each individual’s body mass and taking dorsal photographs to document the recovery process. Wound healing was described macroscopically, as no histological sections were performed.

The recovery followed typical stages of re-epithelialization, tissue regeneration, and pigment cell migration. During the first four days, a thin epithelial layer formed over the wound surface, progressively covering the injured area. This epithelial regeneration involves activation of the skin’s germinative layer, which is essential for cell renewal and wound protection. By day 21, wounds were fully healed, characterized by contraction of the wound edges and the migration of melanophores into the affected area, partially restoring the frogs’ characteristic pigmentation.

### Biopsy does not have a significant effect on body mass

We used body mass as a proxy for health and to evaluate whether the biopsy procedure affected the frogs’ body condition. Results for *G. riobambae* were consistent across laboratory and semi-natural enclosure datasets, as well as when these datasets were analyzed jointly; therefore, we report results from the combined dataset for this species. For visualization purposes, however, trajectories of body mass over time and comparisons of final body mass between control and biopsy groups (Fig. 3) for *G. riobambae* are presented separately for laboratory and semi-natural enclosures. Welch’s t-tests revealed no statistically significant differences in final body mass between biopsy and control groups in any of the sampled species (all *p* > 0.05), suggesting that the biopsy procedure did not negatively affect body condition or recovery by the end of the experiment.

Mean differences in body mass between biopsy and control groups were small across species, ranging from −0.20 g in *Atelopus balios* (with control individuals slightly heavier) to +1.26 g in *Gastrotheca riobambae* (with biopsied individuals slightly heavier). In all species, 95% confidence intervals overlapped zero at both the beginning (May 1) and end (June 1) of the experiment, and group comparisons were not statistically significant at either time point, indicating that relative differences between groups were maintained through time and were not influenced by the biopsy procedure.

To further assess whether the biopsy procedure influenced body mass trajectories, we fitted a linear mixed-effects ANCOVA that included treatment group (biopsy vs. control) and day as fixed effects, and included individual identity as a random intercept. This analysis was conducted for each species separately as body mass varied distinctively between the sampled species. Initial body mass was included as a covariate to control for baseline differences and was a strong predictor of subsequent measurements (β = 0.71 – 0.99), indicating that baseline differences among individuals largely explained variation in mass throughout the experiment. Importantly, treatment group did not have a significant effect on body mass trajectories in any species (all |β| ≤ 0.08; all p ≥ 0.15), indicating that frogs subjected to biopsies maintained body mass comparable to control individuals. Sampling day captured small temporal shifts in body mass (|β| ≤ ∼0.06 g), which were minor relative to baseline differences among individuals and overall body mass. In summary, the groups of frogs subjected to the biopsy procedure are not different from the control group in any of the analyses conducted, with both groups exhibiting similar patterns of body mass fluctuation from the beginning of the experiment (May 1) through the completion of recovery and the end of the study period (June 1). All measurements are available in the Supplementary Material Table 1.

### Post-biopsy reactions

Three types of post-biopsy reactions were observed among specimens. On May 2, one individual of *A. longirostris* (AQSP13) developed a hematoma around the wound characterized by blood accumulation under the skin likely caused by handling. The hematoma gradually reabsorbed over several days without intervention (Fig. 4). On May 5, one specimen of *A. longirostris* (AQSP14) exhibited localized inflammation and pus formation around the wound indicating a possible mild infection or immune response. This condition resolved spontaneously as the inflammation subsided and the purulent secretion disappeared. Finally, one individual of *A. balios* (AQSP05) showed a whitish secretion at the wound site (Fig. 4), interpreted as a regenerative response associated with the accumulation of dead cells, which also resolved without further issues. Monitoring continued for several additional days, documenting melanophore migration into the scarred area. This progressive repigmentation rendered the wounds nearly indistinguishable in some specimens (Figs. 2, 4), restoring the uniformity of skin coloration.

**Figure 2.**
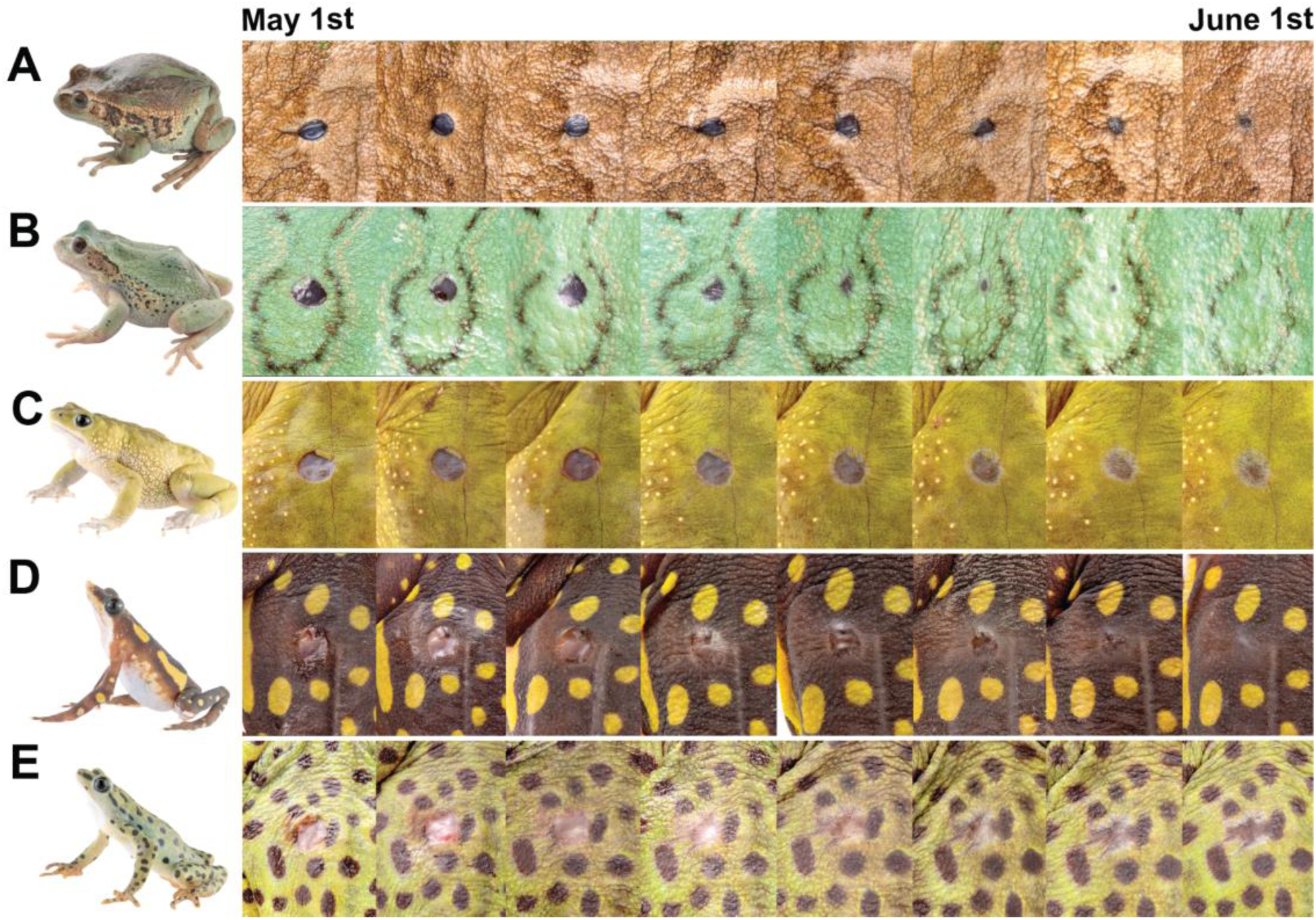
Post-biopsy wound healing across sampled species. Sequential images taken every three days from May 1 to June 1, 2024, document recovery following skin biopsy, including re-epithelialization and repigmentation of the biopsy site. **A.** *Gastrotheca riobambae* (semi-enclosed conditions) Images: Luis A. Coloma. **B.** *G. riobambae* (laboratory conditions). Images: Steven Guevara S. **C.** *Atelopus bomolochos*. Images: Amanda B. Quezada. **D.** *Atelopus longirostris*. Image: José Vieira. **E.** *Atelopus balios*. Images: Amanda B. Quezada. Images are ordered chronologically from left to right.

**Figure 3.**
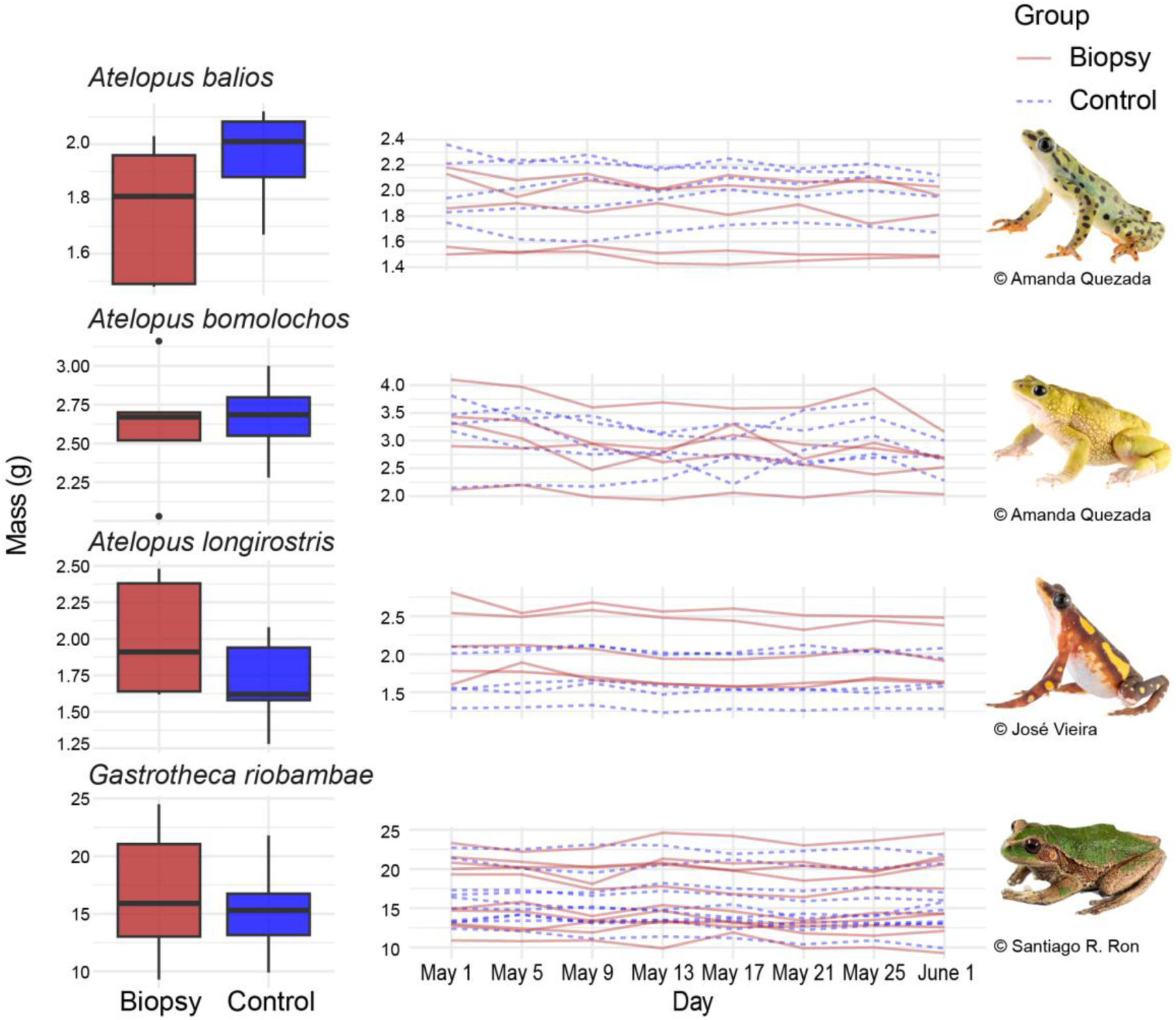
Body mass variation over the course of the experiment as an indicator of body condition. On the left panel, boxplots show a comparison of body mass at the end of the experiment between biopsy (red) and control (blue) groups for each species. Welch’s t-tests revealed no statistically significant differences in final body mass between biopsy and control groups in any of the sampled species (all *p* > 0.05). On the right panel, time series plots show individual body mass trajectories from May 1 to June 1, 2024, the period during which wound healing progressed to full closure and re-epithelialization. Body mass was measured every three days. Across all species, body mass at both the beginning and end of the experiment did not differ between biopsy (solid red lines) and control individuals (dashed blue lines). Observed fluctuations over time likely reflect normal variation associated with hydration status and food intake rather than effects of the biopsy procedure.

**Figure 4.**
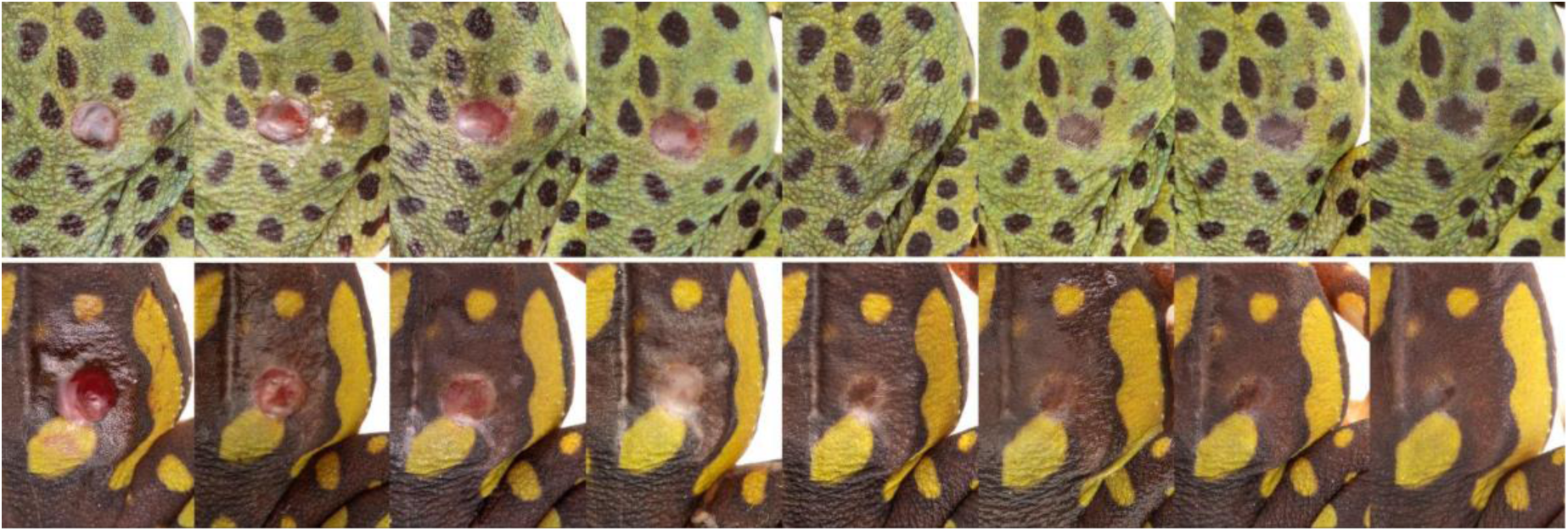
Post-biopsy reactions observed in two individuals. The upper panel shows an individual of *A. balios* (AQSP05) exhibiting a whitish secretion at the biopsy site. The lower panel shows an individual of *A. longirostris* (AQSP13) that developed a localized hematoma around the biopsy site, characterized by subcutaneous blood accumulation likely associated with handling during sampling. Neither of these reactions resulted in mortality, and both resolved without veterinary treatment.

### Field-settings recovery

We were able to anecdotally document post-biopsy wound recovery in free-living individuals through ongoing field monitoring of three *Atelopus* species. Photographs taken during these encounters show progressive re-epithelialization of the biopsy site, evident as the formation of a pale, translucent tissue layer covering the wound. Although complete closure was not observed in the final photographs for any individual, this appearance corresponded to the final stages of healing (approximately the week prior to full closure) in lab-bred individuals. The longer time required to reach a comparable stage in free-living frogs suggests that wound healing under natural field conditions may proceed more slowly than under controlled laboratory settings. In all documented cases, biopsy sites appeared reduced in size by the time of the first post-biopsy encounter, which occurred 8 days after sampling for *A. coynei*, 10 days for *A.* sp. (*aff. longirostris*), and 13 days for *A. laetissimus*. These observations revealed visible tissue regeneration with no signs of infection, necrosis, or abnormal inflammation, consistent with normal healing trajectories observed in captive individuals. For all three individuals, the final photographs showed full re-epithelialization of the wound, although repigmentation and complete closure were not yet evident at the time of observation—approximately one month post-procedure for *A. laetissimus*, and 14 and 19 days post-biopsy for *A. coynei* and *A.* sp. (*aff. longirostris*), respectively.

## DISCUSSION

Our study provides evidence that small (2-mm) non-lethal skin biopsies are a safe and effective method for obtaining amphibian tissue samples. In the lab, for several species of *Atelopus* (Bufonidae) and one species of *Gastrotheca* (Hemiphractidae), wounds fully healed within three weeks and showed signs of melanophore migration leading to tissue repigmentation. These findings are further supported by field observations in three *Atelopus* species, in which individuals previously biopsied and released were later encountered during independent surveys, allowing us to document post-biopsy recovery under natural conditions. In these field-sampled individuals, the healing process closely resembled the stages of re-epithelialization observed in laboratory-kept frogs, indicating that wound healing reliably occurs in natural environments (Figs. 1, 2). However, recovery in the field appeared to progress more slowly than under laboratory conditions, with a lag of approximately one to two weeks, suggesting that environmental factors may influence the rate of wound healing (Figs. 1, 2). Given the limited number of field-sampled individuals and the opportunistic nature of these observations, we cannot rule out the possibility of higher post-biopsy mortality or subtle effects on body condition under field conditions, although neither were detected under lab conditions.

Additionally, no biopsy-related mortality was observed during this study, and body mass in biopsied individuals remained stable relative to both control animals and initial measurements (Fig. 3). It is worth noting, however, that outside the scope of this study, we have observed three cases of mortality associated with the biopsy procedure, out of a total of 173 samples collected using this method (Navarrete pers. comm.). In these cases, wounds showed no signs of infection and no overt clinical symptoms were observed, suggesting that increased stress associated with handling and/or sampling may have contributed to mortality in these individuals, or that other undetermined causes were involved. While these instances represent a very low proportion (1.7%) of the total individuals sampled, they highlight that a level of risk is inherent to any invasive sampling technique, including handling (Waddle et al. 2008) or toe clipping. Again, we note that there were two mortality events of control animals in the lab experiment (2 of 25 total control individuals), further emphasizing the potential effects of handling stress even when procedures are conducted by highly trained and experienced personnel. In addition, as with other non-lethal methods, outcomes can be influenced by factors such as individual condition, species-specific sensitivity and behavior, and handling stress (Phillott et al. 2007; Grafe et al. 2011), underscoring the importance of proper training and post-sampling monitoring.

The rapid recovery we observed is consistent with the high regenerative capacity of amphibian skin, which possesses a robust epithelial renewal system and specialized immune defenses that promote healing (Godwin and Rosenthal 2014). The sequential pattern of re-epithelialization, melanophore migration, and repigmentation mirrors what has been reported in other amphibians subjected to minor dermal injuries without the formation of a remarkable scar (Ishii et al. 2021; Godwin and Rosenthal 2014). Importantly, the timeline of recovery was also comparable to that reported for other amphibian species. Across all three contexts examined—laboratory conditions, outdoor seminatural enclosures, and field observations—re-epithelialization, wound closure, and repigmentation were nearly complete by 30 days post-biopsy, a timescale comparable to that reported for other amphibian species, such as *Xenopus laevis,* in which wound healing following biopsy was nearly complete by 28 days post-injury (Garvey Griffith et al. 2025),, and the newt *Cynops pyrrhogaster*, which exhibits comparable recovery stages between approximately 15 days (stage 4) and two months (stage 5) post-injury (Ishii et al. 2021). Occasional minor complications, such as hematoma, localized inflammation, or transient exudate, were rare and resolved without intervention, suggesting that standard hygienic precautions and proper handling are sufficient to minimize the risk of infection.

Our results indicate that the biopsy procedure did not adversely affect body condition in the lab. Neither final body mass nor body mass trajectories over time differed significantly between biopsy and control groups, suggesting that the small incision was unlikely to have substantially interfered with feeding, hydration, or locomotion, and that any sampling-associated stress did not produce detectable, longer-term effects on body condition during the wound-healing period. Although body mass fluctuated throughout the experiment, these trajectories were similar between groups and likely reflected normal day-to-day variation associated with food intake and hydration rather than effects of the biopsy procedure.

The lack of significant differences between animals maintained in laboratory and outdoor semi-natural enclosures further supports the robustness of the technique for wild populations. Although controlled environments facilitated daily monitoring, the consistent recovery of *G. riobambae* under outdoor conditions suggests that similar outcomes can be expected in the field when individuals are released after sampling. Further, field observations of different *Atelopus* species in Ecuador and Colombia likewise confirmed survival and visible wound closure post-release, underscoring the potential for applying this method during in-situ conservation efforts. While promising, some limitations warrant consideration. The sample sizes were modest, and long-term monitoring beyond one month was not conducted. Histological assessment of tissue regeneration could strengthen the evaluation of complete healing. Additionally, the control deaths in our experiment highlight the sensitivity of amphibians to changes in housing conditions and to handling, reinforcing the need for careful handling and husbandry when performing experimental manipulations. Future studies incorporating larger sample sizes, additional taxa, and extended follow-up periods will help refine best practices and assess whether species-specific factors such as body size (snout-vent length) or skin thickness influence recovery rates.

The outcomes we report for the skin biopsy technique are broadly comparable to those described for more widely used methods such as toe-clipping. A recent meta-analysis found that the majority of studies evaluating the effects of toe-clipping in frogs and salamanders reported no strong evidence of detrimental impacts (Zemanova et al. 2025). For example, a laboratory study on leopard frogs (*Rana pipiens*) found no differences in survival rates between toe-clipped individuals (2, 3, 4, 8, or 12 toes clipped, n = 10 for each group) and control frogs (n = 50), nor in the number of days individuals survived over a 13-week period (Ginnan et al. 2014)—a pattern generally consistent with our observations during the post-biopsy recovery period. However, that same meta-analysis also highlighted several welfare-related effects associated with toe-clipping, including impaired mobility immediately following digit removal, elevated urinary corticosterone levels, and reduced daily weight gain, indicating that subtle or transient costs may occur even when overt negative outcomes are not detected.

Inflammation represents a response common to both sampling techniques, having been reported following toe-clipping in some individuals (Golay and Durrer 1994; Phillott et al. 2011) as well as observed following skin biopsy in this study. However, toe-clipping has also been associated with additional risks—including infection, decreased return rates (often interpreted as a proxy for increased mortality, particularly when multiple digits are removed; (Parris and McCarthy 2001; Olivera-Tlahuel et al. 2017), and abnormal digit regeneration (Stock and Bryant 1981)—that we did not observe following skin biopsy. This contrast should be interpreted cautiously, as toe-clipping has been applied far more extensively across studies, encompassing larger sample sizes, longer monitoring periods, and broader taxonomic representation than captured in our work. Consequently, some risks associated with skin biopsy may remain undetected, emphasizing the need for standardized experimental designs and formal power analyses when evaluating non-lethal sampling techniques (Zemanova et al. 2025).

Beyond reducing individual-level impacts, the biopsy method offers clear analytical advantages over other non-lethal sampling approaches by providing sufficient tissue for a wide range of downstream applications. These include chemical characterization of amphibian skin, quantification of toxins such as tetrodotoxin (TTX), skin transcriptomics and proteomics, morphological analyses, disease detection, in vitro skin culturing, and microbiome characterization at both compositional and functional levels. For chemical analysis in particular—one of the primary purposes for which we have used this technique—skin biopsies offer several advantages over skin swabs and other methods such as the Transcutaneous Amphibian Stimulator (TAS). Because the diameter of the sampled tissue is known, biopsies allow more accurate estimation of whole-individual toxicity (Hanifin et al. 1999; Bucciarelli et al. 2014; Vaelli et al. 2020) than skin swabs or bath-based methods, which depend on active secretion and often result in diluted concentrations. In addition, based on our experience, skin biopsies can provide sufficient, high-quality RNA for transcriptomics, enabling a broad range of downstream analyses directly from skin tissue. While other non-lethal methods, such as toe clipping, have been used for targeted RNA-based applications (e.g., virus detection; (Parry et al. 2023), the limited amount of tissue obtained may constrain their utility for broader transcriptomic studies.

Notwithstanding the methodological advantages associated with skin biopsies, this approach is not intended to replace whole-animal collections, which remain essential for many research objectives, biodiversity documentation, and conservation (Nachman et al. 2023). Whole specimens provide irreplaceable value for detailed anatomical and morphological studies, comprehensive physiological and developmental analyses, and the establishment of museum voucher specimens that serve as permanent records of biodiversity, and allow for comparative studies across time and space (Suarez and Tsutsui 2004; Meineke et al. 2018). Rather, biopsy-based approaches provide a complementary tool for situations in which whole-animal collection is impractical, ethically constrained, or incompatible with conservation goals. This distinction is particularly relevant for *Atelopus* and other highly threatened amphibians, where populations are often small, fragmented, and declining (Lötters et al. 2023). In such cases, population-level inference may require sampling multiple individuals while minimizing impacts on population viability, making non-lethal approaches necessary rather than optional. For reference, we present a comparison between commonly used sampling methods and skin biopsy in terms of invasiveness, repeatability, types of data obtained, ethical considerations, and limitations across species in Table 1.

**Table 1.** Comparison of commonly used sampling methods, including relative invasiveness, benefits and data obtained, limitations, and ethical considerations.

|  | Toe clipping | Skin biopsy | Skin/buccal swabbing | Whole animal collection |
| --- | --- | --- | --- | --- |
| **Level of invasiveness** | Moderate | Moderate | Low | High |
| **Data obtained** | Low–moderate: DNA; individual identification; disease detection | Moderate–high: DNA; toxins (e.g. TTX); gene expression; microbiome data; disease detection; skin structure | Low–moderate: DNA; microbiome data; some metabolites/toxins; disease detection | Very high: Full anatomy, physiology, chemistry, genomics, microbiome data; multiple tissue transcriptomics; diet; disease detection and parasite load; voucher as museum specimen; |
| **Recovery time** | Days–weeks | Weeks | Immediate | N/A |
| **Repeatability over time** | Low (cumulative impacts) | Moderate (after sufficient recovery) | High (frequent repetition possible) | None |
| **Ethical /**<br>**conservation**<br>**implications** | Potential locomotor<br>and fitness effects;<br>can cause death | Short-term injury;<br>requires careful site<br>selection; can cause<br>death | Minimal stress and<br>injury | Irreversible loss of<br>individuals |
| **Key**<br>**limitations** | Limited tissue;<br>behavioral effects | Not feasible for very<br>small individuals;<br>expertise required | Low biomass;<br>surface-limited<br>data | Not repeatable; poses<br>substantial risk to<br>already declining<br>populations |

In conclusion, the 2-mm skin biopsy technique represents a practical compromise between scientific rigor and animal welfare. By enabling high-quality tissue collection while preserving individual survival, this approach is especially well suited for research on species of conservation concern, likely even beyond amphibians. As amphibian populations continue to decline globally, the development and careful validation of non-lethal sampling methods will be essential to ensure that research efforts do not become an additional source of risk for the species it seeks to protect.

## Acknowledgments

We thank J. Vieira and S. S. Amini for their assistance in developing and refining the biopsy protocol. We also thank K. Velez and T. Hildwein for kindly providing photographs documenting post-biopsy recovery under field conditions. We are grateful to the staff at Centro Jambatu for performing animal husbandry and care for the individuals used in this study. We also thank Ammon Corl for helpful comments that improved the manuscript. Funding was provided by NIH NIGMS R35GM150574 to RDT and by fellowships from the American Association of University Women and Philomathia to MJN. The Ecuadorian Ministry of Environment provided permits for specimen collection and genetic research: Nro. MAE-DNB-CM-2022-0261.

